# Constrained G4 structures unveil topology specificity of known and new G4 binding proteins

**DOI:** 10.1101/2021.04.06.438633

**Authors:** A. Pipier, A. Devaux, T. Lavergne, A. Adrait, Y. Couté, S. Britton, P. Calsou, J.F. Riou, E. Defrancq, D. Gomez

## Abstract

G-quadruplexes (G4) are non-canonical secondary structures consisting in stacked tetrads of hydrogen-bonded guanines bases. An essential feature of G4 is their intrinsic polymorphic nature, which is characterized by the equilibrium between several conformations (also called topologies) and the presence of different types of loops with variable lengths. In cells, G4 functions rely on protein or enzymatic factors that recognize and promote or resolve these structures. In order to characterize new G4-dependent mechanisms, extensive researches aimed at identifying new G4 binding proteins. Using G-rich single-stranded oligonucleotides that adopt non-controlled G4 conformations, a large number of G4-binding proteins have been identified *in vitro*, but their specificity towards G4 topology remained unknown.

Constrained G4 structures are biomolecular objects based on the use of a rigid cyclic peptide scaffold as a template for directing the intramolecular assembly of the anchored oligonucleotides into a single and stabilized G4 topology. Here, using various constrained RNA or DNA G4 as baits in human cell extracts, we establish the topology preference of several well-known G4-interacting factors. Moreover, we identify new G4-interacting proteins such as the NELF complex involved in the RNA-Pol II pausing mechanism, and we show that it impacts the clastogenic effect of the G4-ligand pyridostatin.

## Introduction

In the last twenty years G-quadruplex structures (G4) emerged as cis-acting factors impacting almost all DNA and RNA transactions. G4 in telomeric sequences were first shown to play essential roles in telomeres capping and telomeres replication by telomerase ^1^. Now, G4 are associated with the firing of DNA replication origins ^2, 3^, transcription initiation and termination, mRNA processing, mRNA transport ^4–6^, translation ^7^ and mitochondrial maintenance ^8^.

G4 are noncanonical secondary structures formed by stacked tetrads of Hoogsteen hydrogen-bonded guanines bases, which are stabilized through the coordination of physiologically relevant cations (Na^+^, K^+^). G4 can result from the intramolecular folding of a unique G-rich sequence or from the intermolecular assembly of different G-rich containing strands ^9^. An essential feature of G4 is their intrinsic polymorphic nature: numerous *in vitro* studies have revealed their ability to adopt different conformations, also called topologies ^10^. Indeed, depending on the length and the composition of the sequence, as well as the environmental conditions (including the nature and concentration of metal cations, and local molecular crowding), a G4-forming sequence can adopt different topologies, in which the strands are in parallel, antiparallel or hybrid conformations, with the co-existence of different types of loops (lateral, diagonal or propeller) of variable lengths ^9–11^. In particular, this polymorphism is exacerbated for the human telomeric sequence and leads to intricate structural mixtures ^12^.

In cells, the impact of G4 on cellular metabolism is mainly associated with protein or enzymatic factors that bind, stabilize or resolve these structures. The folding of G-rich sequences into a G4, on DNA and RNA molecules, is associated with the formation of DSBs, transcription and translation repression and the alteration of the RNA processing ^13–15^. To handle these major threats, cells use a battery of DNA and RNA helicases to control G4 formation ^16, 17^. Notably, most of helicases resolving DNA G4 are associated, when mutated, with genetic disorders, progeria and cancer progression (WRN, BLM, FANCJ, RTEL), underlying the major impact of G4 structures on cell fitness ^18, 19^. In addition to helicases, the formation of G4 structures in cells is counteracted by proteins that bind single-stranded nucleic acids ^19, 20^ through their OB-fold, RRM or RGG interacting motifs ^21–23^. Interestingly, RGG motif containing proteins also promote G4 stabilization ^24^ and control mRNAs localization through interacting with G4 ^25^. A major impact of G4 structures in cells is related to transcription ^4^. Found enriched on promoters and transcriptional start sites (TSS) ^26, 27^, G4 structures have been shown to act predominantly as transcriptional repressors ^4, 13, 15^, although some G4 have been also described as involved in transcription activation ^28, 29^. Furthermore, the presence of G4 motifs in the TSS proximal regions is associated with RNA-Pol II pausing sites and R-loops formation, two different factors promoting RNA-Pol II arrests and transcription-dependent DNA breaks ^30–34^.

Given the increasing roles of G4 structures in cellular metabolism, extensive researches have been conducted in the last years in order to identify new G4-dependent mechanisms. Notably classical pull-down approaches identified hundreds of proteins associated to G-rich oligonucleotides forming G4 structures ^35–40^. In solution, G-rich single-stranded molecules are in equilibrium between unfolded and folded states, and thus numerous identified G4 binding proteins are also able to recognize unfolded G-rich sequences ^20^. In addition, G4 derived from single-stranded oligonucleotides can adopt different topologies ^9, 11, 41^, that precludes to establish the specific contribution of each G4 topology to protein binding.

In this context, we have developed an approach to constrain the G4 into a single well-defined topology. The strategy is based on the use of a rigid cyclic peptide scaffold as a template for directing the intramolecular assembly of the anchored RNA or DNA oligonucleotides ^42–45^. Moreover, such locked G4 display a thermal stability significantly higher than unconstrained G4 that strongly reduces the possibility to form unfolded single-stranded sequences. These constrained systems represent original tools, that we have used here for the identification and characterization of proteins interacting with a well-defined RNA or DNA G4 topology. In this study we identified through affinity purifications coupled to mass spectrometry (MS)-based quantitative proteomics a set of human proteins associated to locked G4 structures. Notably, this approach allowed us to identify NELF proteins as a new G4-interacting complex, leading us to investigate the impact of RNA-Pol II pausing mechanism into the response to G4 stabilization by G4 ligands.

## Results

### Identification and characterization of G4 associated proteins

To identify human proteins interacting with G4 structures we performed classical pull-down assays followed by MS-based quantitative proteomic analysis. Various constrained G4 topologies based on the telomeric sequence (excepted for **5**) were used (Supplementary Figure S1): systems **1, 6** and **7** depict a parallel topology, systems **2-5** have an antiparallel topology. In our approach, biotin-functionalized G4-constrained molecules **1-7** and the biotin-functionalized duplex-DNA control **8**, were incubated individually with a semi-total human protein extracts prepared from HeLa cells ^46^, before being trapped using streptavidin magnetics beads to isolate interacting proteins (Figure 1B). In a first-round assay, and in order to validate our approach, western-blotting analyses were performed to test the interaction and the binding specificity of some well-established and depicted G4-binding proteins to constrained G4 constructions. From these analyses we observed that eIF4G, WRN, Nucleolin, Mre11, DHX36, hnRNP A1 and CNBP, all well-known G4-interacting proteins ^20^, were enriched using constrained G4 structures compared to the duplex control **8**. Conversely, the KU heterodimer, one of the most abundant human duplex-DNA-binding proteins ^47^, was found enriched using duplex control **8** but was barely detectable on constrained G4 structures (Supplementary Figure S2A-B). Comparative analysis of the G4-interacting proteins enrichment on the six different constrained G4 structures **1a-3 and 5-7** shows a differential binding for these human proteins. Indeed, while eIF4G and Mre11 proteins are particularly enriched on constrained G4 **2** and **3** (*i.e.* antiparallel topology with two lateral loops), DHX36 and hnRNP A1 proteins are abundant on constrained G4 **1a, 6** and **7** (*i.e.* parallel topology without loops) (Supplementary Figure 2A-B). Altogether, these first assays confirm that our pull-down strategy using constrained G4 structures allowed to both identify G4 binding proteins and to discriminate important aspects of their structural interaction with G4. Thus, our approach represents a powerful tool to find new proteins recognizing particular G4 conformations and prompted us to proceed to an extensive MS-based quantitative proteomic analysis of human proteins interacting with two particulars constrained G4 structures, systems **1a** (*i.e.* parallel without loops) and **2** (*i.e.* antiparallel with two lateral loops), compared to control **8** (Figure 1A-B).

**Figure 1:**
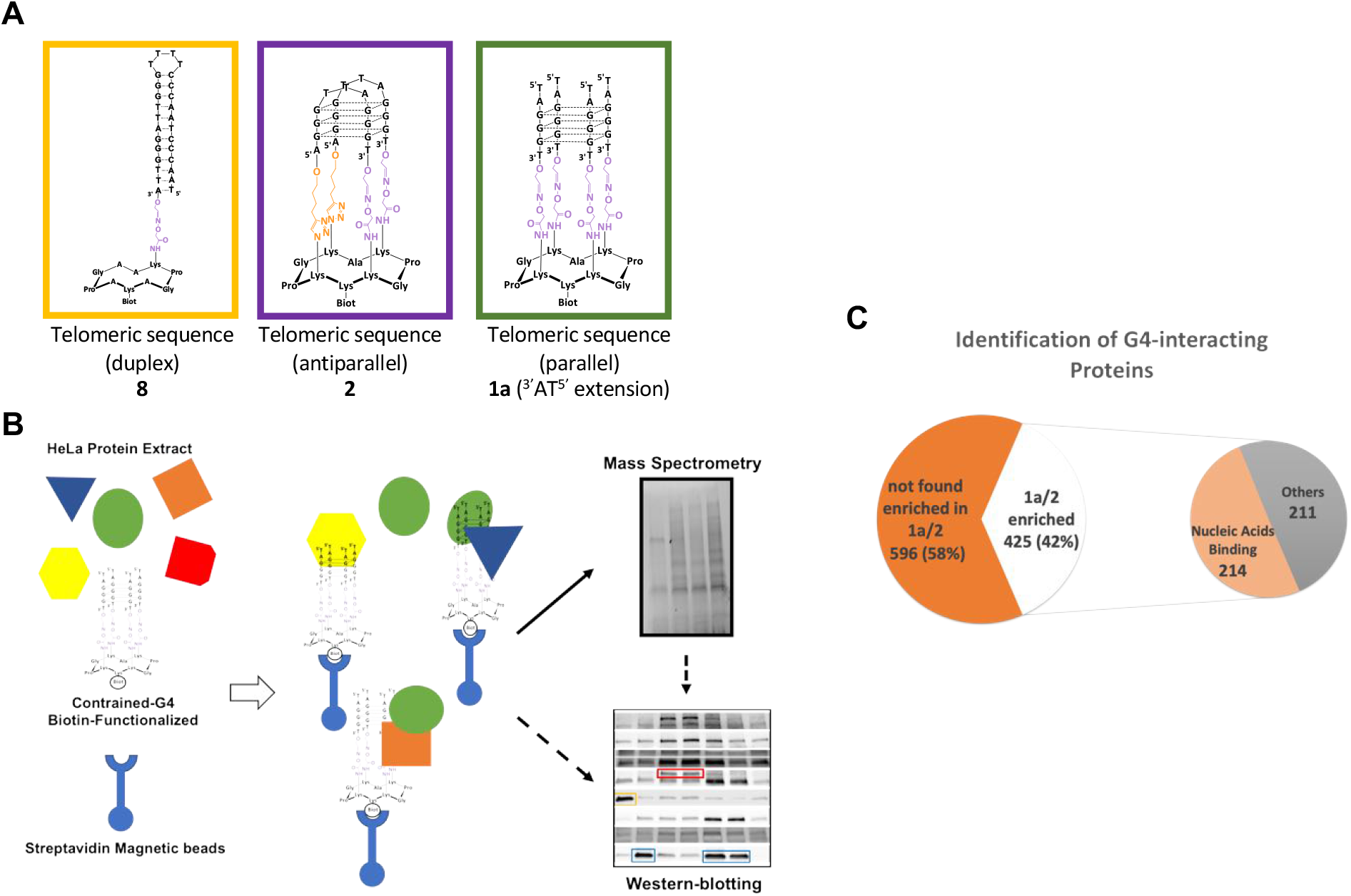
**A** Schematic representation of constrained DNA structures used in the pull-down assay. **B** Global strategy to identify constrained G4 interacting proteins from human cells. Biotin-functionalized G4-constrained molecules (**1a** and **2**) and the biotin-functionalized duplex-DNA control **8** were individually mixed with a semi-total human protein extracts from HeLa cells, then trapped by streptavidin magnetics beads to isolate interacting proteins. Protein identification was obtained from MS-based quantitative proteomic analysis and further characterized by western-blotting (arrows), or directly by western-blotting (dashed arrow). **C** Diagram showing the differential enrichment of human proteins on constrained G4 structures relative to control duplex DNA. G4 enriched proteins refer to proteins found enriched on **1a** and/or **2** G4 constructions relative to the duplex control **8**. 214 out of 425 proteins found enriched on constrained G4 have been shown to interact with nucleic acids. Differentially interacting proteins were sorted out using a fold change ≥ 2 and p-value < 0.05, allowing to reach a false discovery rate (FDR) inferior to 5% according to the Benjamini-Hochberg procedure.

Based on three independent experiments, MS-based proteomic analyses identified a total of 1021 proteins interacting with constrained structures (see Materials and Methods for identification conditions) (Supplementary Table 1). This total corresponds to the sum of proteins interacting with **1a**, **2** and control **8** (duplex) (Figure 1C and Supplementary Table 1). Using a label-free quantification and statistical filtering to compare the abundances of the proteins eluted from different constructions (see Materials and Methods for filter conditions), we identified 425 proteins enriched on constrained G4 structures **1a-2** compared to duplex control **8** (fold change ≥ 2 and p-value < 0.05, allowing to reach a false discovery rate (FDR) inferior to 5%) (Figure 1C and Supplementary Table 2). These proteins belong to six significant enriched KEGG pathways cluster (p < 0.05): Spliceosome, RNA transport, RNA degradation, mRNA surveillance, DNA replication and Homologous Recombination (Figure 2A). To go further, enriched GO Biological Processes and Molecular Functions terms were also determined. This analysis revealed that proteins enriched on contrained-G4 structures are mainly associated with DNA and RNA transactions (Figure 2B-C), in agreement with current knowledge on genomic localization and biological function of G4. In line with these data, 214 out of 425 proteins enriched on G4 structures have been described as nucleic acid interacting factors, as indicated by terms from GO Molecular functions data analyses (Figure 1C and Supplementary Table 3). Furthermore, the KEGG pathways clusters from these 214 nucleic acid binding proteins correspond to almost the same biological processes defined by the complete set of G4-interacting proteins (Supplementary Figure S3). An additional analysis of nucleic acid binding proteins enriched on constrained G4 structures defines five functional groups covering (i) ATP dependent DNA/RNA helicases activities, (ii) hnRNP proteins, (iii) proteins involved in the polyadenylation process, (iv) a large group of proteins involved in splicing and (v) proteins related to small nuclear ribonucleoprotein complexes (Table 1).

**Figure 2:**
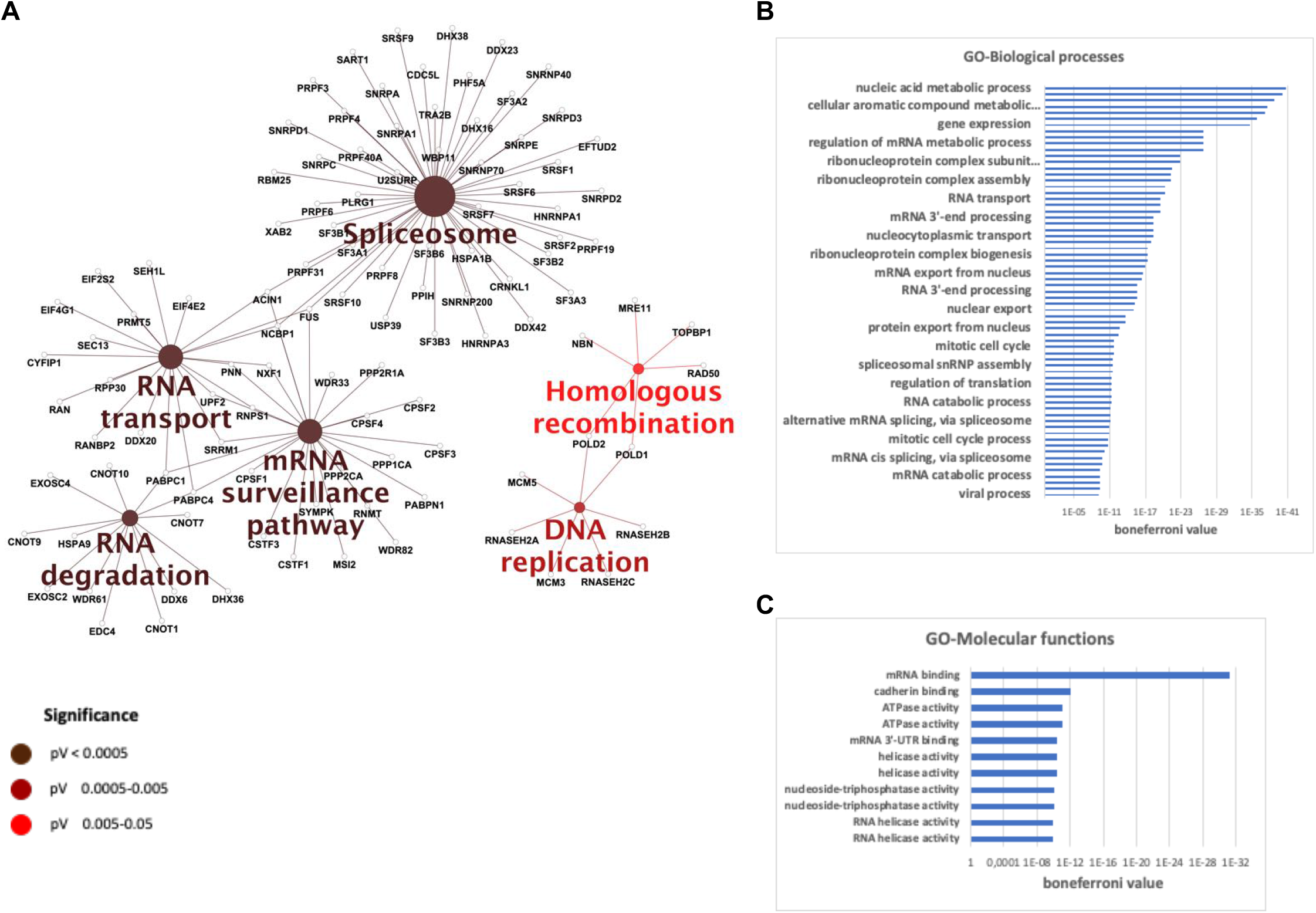
Most significant pathways and processes covered by constrained G4 interacting proteins. (**A**) Enriched KEGG pathways. Gene Ontology terms, (**B**) GO-Biological processes and (**C**) GO-Molecular Functions for the 425 proteins found enriched on constrained G4 structures. A rightsided (Enrichment) test based on the hyper-geometric distribution was performed on the corresponding Entrez gene IDs for each gene list and the Bonferroni adjustment (p<0.05).

**Table 1:** Manually curated functional groups from nucleic acid binding proteins enriched on constrained G4 structures

| Mapped IDs | Gene Name | Mapped IDs | Gene Name |
| --- | --- | --- | --- |
| **DDX1** | ATP-dependent RNA helicase DDX1 | **SF1** | Splicing factor 1 |
| **DDX20** | Probable ATP-dependent RNA helicase DDX20 | **SF3A1** | Splicing factor 3A subunit 1 |
| **DDX23** | Probable ATP-dependent RNA helicase DDX23 | **SF3A2** | Splicing factor 3A subunit 2 |
| **DDX3X** | ATP-dependent RNA helicase DDX3X | **SF3A3** | Splicing factor 3A subunit 3 |
| **DDX41** | Probable ATP-dependent RNA helicase DDX41 | **SF3B1** | Splicing factor 3B subunit 1 |
| **DDX42** | ATP-dependent RNA helicase DDX42 | **SF3B2** | Splicing factor 3B subunit 2 |
| **DDX52** | Probable ATP-dependent RNA helicase DDX52 | **SF3B3** | Splicing factor 3B subunit 3 |
| **DDX6** | Probable ATP-dependent RNA helicase DDX6 | **SF3B6** | Splicing factor 3B subunit 6 |
| **DHX16** | Pre-mRNA-splicing factor ATP-dependent RNA helicase DHX16 | **SRSF1** | Serine/arginine-rich splicing factor 1 |
| **DHX29** | ATP-dependent RNA helicase DHX29 | **SRSF10** | Serine/arginine-rich splicing factor 10 |
| **DHX30** | Putative ATP-dependent RNA helicase DHX30 | **SRSF11** | Serine/arginine-rich splicing factor 11 |
| **DHX36** | ATP-dependent RNA helicase DHX36 | **SRSF2** | Serine/arginine-rich splicing factor 2 |
| **DHX38** | Pre-mRNA-splicing factor ATP-dependent RNA helicase DHX38 | **SRSF6** | Serine/arginine-rich splicing factor 6 |
| **DHX40** | Probable ATP-dependent RNA helicase DHX40 | **SRSF7** | Serine/arginine-rich splicing factor 7 |
|  |  | **SRSF9** | Serine/arginine-rich splicing factor 9 |
| **hnRNP A1** | Heterogeneous nuclear ribonucleoprotein A1 |  |  |
| **hnRNP A2B1** | Heterogeneous nuclear ribonucleoproteins A2-B1 | **SNRNP200** | U5 small nuclear ribonucleoprotein 200 kDa helicase |
| **hnRNP A3** | Heterogeneous nuclear ribonucleoprotein A3 | **SNRNP40** | U5 small nuclear ribonucleoprotein 40 kDa protein |
| **hnRNP F** | Heterogeneous nuclear ribonucleoprotein F | **SNRNP70** | U1 small nuclear ribonucleoprotein 70 kDa |
| **hnRNP H1** | Heterogeneous nuclear ribonucleoprotein H1 | **SNRPA** | U1 small nuclear ribonucleoprotein A |
| **hnRNP H3** | Heterogeneous nuclear ribonucleoprotein H3 | **SNRPA1** | U2 small nuclear ribonucleoprotein A |
| **hnRNP L** | Heterogeneous nuclear ribonucleoprotein L | **SNRPC** | U1 small nuclear ribonucleoprotein C |
| **hnRNP R** | Heterogeneous nuclear ribonucleoprotein R | **SNRPD1** | Small nuclear ribonucleoprotein Sm D1 |
|  |  | **SNRPD2,SNRPD1** | Small nuclear ribonucleoprotein Sm D2 |
| **CPSF1** | Cleavage and polyadenylation specificity factor subunit 1 | **SNRPD3** | Small nuclear ribonucleoprotein Sm D3 |
| **CPSF2** | Cleavage and polyadenylation specificity factor subunit 2 | **SNRPE** | Small nuclear ribonucleoprotein E |
| **CPSF3** | Cleavage and polyadenylation specificity factor subunit 3 | **SNRPN** | Small nuclear ribonucleoprotein-associated protein N;SNRPN |
| **CPSF4** | Cleavage and polyadenylation specificity factor subunit 4 |  |  |
| **CRNKL1** | Crooked neck-like protein 1 |  |  |
| **CSTF1** | Cleavage stimulation factor subunit 1 |  |  |
| **CSTF3** | Cleavage stimulation factor subunit 3 |  |  |

Next, using UniprotKB, gene Ontology (GO), G4IPDB ^48^ databases and G4 search terms in PubMed, we determined that eighteen proteins found enriched on constrained G4 structures **1a-2** have been already implicated in G4 biology (Table 2).

**Table 2:** Constrained G4 interacting factors found related to G4 on UniprotKB, gene Ontology (GO), PubMed abstract and G4IPDB ^48^ data bases.

| Mapped IDs | Gene Name | PUBMED ID |
| --- | --- | --- |
| ADAR | Double-stranded RNA-specific adenosine deaminase | 24813121, 23381195 |
| CNBP | Cellular nucleic acid-binding protein | 23774591, 28329689, 24594223,<br>26332732, 31219592 |
| DDX1 | ATP-dependent RNA helicase | 29731414 |
| DDX42 | ATP-dependent RNA helicase | 31287417 |
| DHX36 | ATP-dependent RNA helicase | 29269411, 28069994, 25653156,<br>25611385, 24151078, 22238380,<br>21149580, 18842585, 16150737 |
| DNMT1 | DNA (cytosine-5)-methyltransferase 1 | 30275516 |
| EWSR1 | RNA-binding protein | 21244633, 21561087, 22214309 |
| FUS | RNA-binding protein | 18776329, 19749353, 23521792,<br>24251952, 28575444, 29434328,<br>29800261 |
| hnRNP A1 | Heterogeneous nuclear ribonucleoprotein A1 | 9188487, 19282454, 20213319,<br>24371143, 24831962, 26930004,<br>28510424, 29361764, 30247678,<br>31311954 |
| hnRNP A2B1 | Heterogeneous nuclear ribonucleoproteins A2/B1 | 15302914, 17716999 |
| hnRNP A3 | Heterogeneous nuclear ribonucleoprotein A3 | 27623008, 23381195 |
| hnRNP F | Heterogeneous nuclear ribonucleoprotein F | 29269483 |
| hnRNP H1 | Heterogeneous nuclear ribonucleoprotein H | 26930004, 27623008 |
| MID1 | E3 ubiquitin-protein ligase Midline-1 | 21930711 |
| Mre11A (*) | Double-strand break repair protein MRE11 | 16116037 |
| RIF1 | Telomere-associated protein | 26436827, 29348174, 29357064,<br>30510058, 31197198 |
| SF3B3 | Splicing factor 3B subunit 3 | 23381195 |
| SRSF1 | Serine/arginine-rich splicing factor 1 | 24771345 |

We compared our results with two recent studies that identified proteins associated with RNA G4 structures ^35, 36^ and found that 98 out of the 425 proteins identified in our study have been already shown to interact with G4 structures (Figure 3A). Fourteen factors are in common in the three studies, including DHX36 and DDX3 proteins, two major G4 resolvases (Figure 3A). Finally, in order to further investigate the association of constrained G4 interacting proteins identified here with G4 biological functions, we compared our list of 425 proteins with the list of 758 G4 sensitizers genes, the deficiency of which leads to increased sensitivity to G4 ligands, established by Zyner et al ^49^. From this analysis, we determined that at least 62 out of the 425 proteins enriched on constrained G4 structures **1a-2** were reported as G4 sensitiser proteins (Figure 3B).

**Figure 3:**
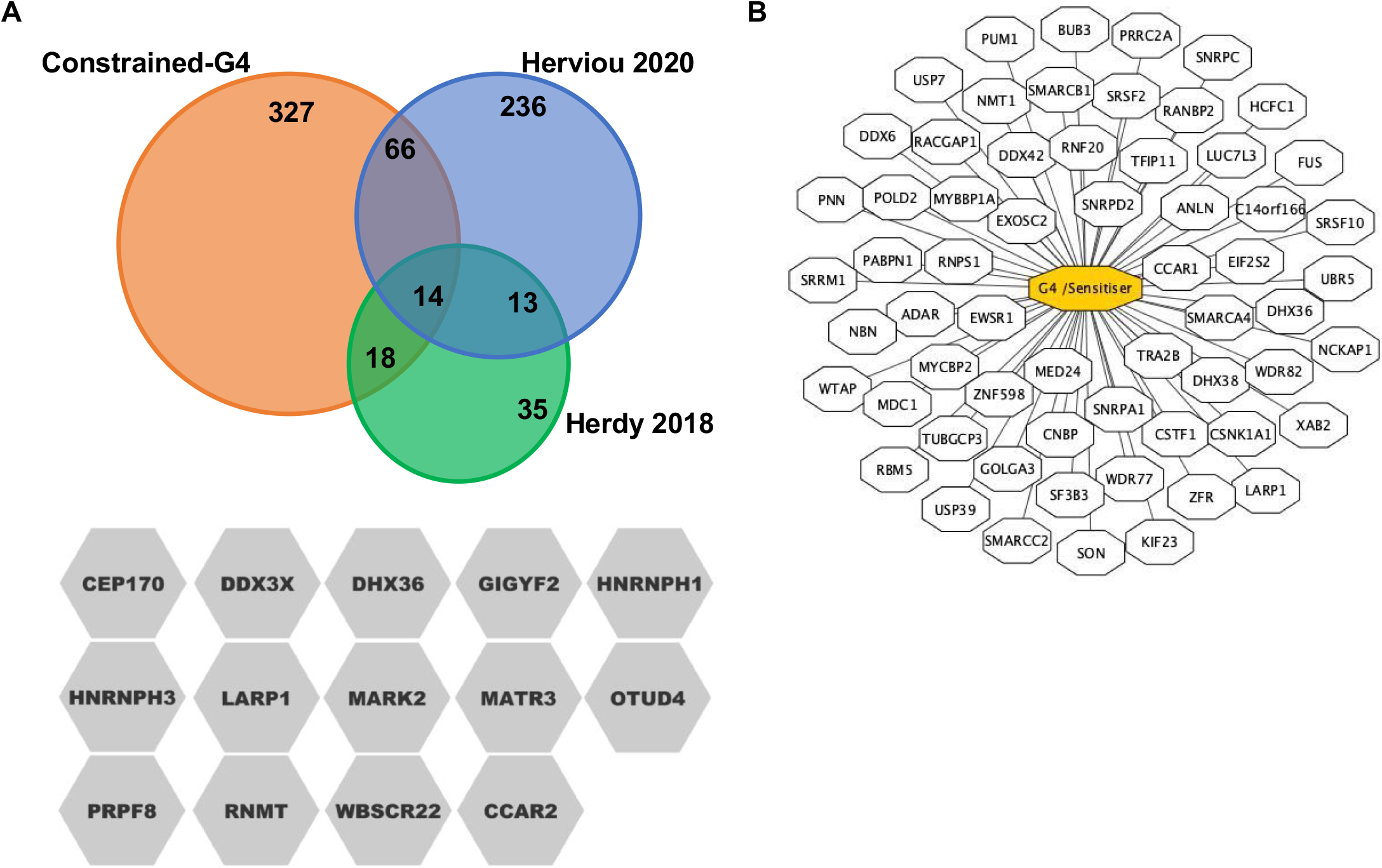
Contrained-G4 interacting factors are associated with RNA-G4 binding activities and with the sensitisation to small molecules that stabilize G4 structures. **A** Venn diagram showing the overlap of our study (orange) with the RNA-G4 interacting proteins identified in Herdy ^35^ (green) and Herviou ^36^. 98 out of 425 proteins identified in our study were known to interact with RNA-G4 structures, with 14 indicated factors common to three studies. **B** Schematic representation of constrained G4 interacting proteins identified in our study that are associated to an increased sensitivity to G4 ligands, established by Zyner et al ^49^. From this analysis we determined that 62 out of 425 proteins were reported as G4 ligands sensitisers.

### Different sets of proteins are enriched on particular G4 conformations

Statistical analysis of the abundances of proteins enriched on G4 structures **1a-2** shows differential interactions for some of them with the two constrained G4 conformations. Indeed, among the proteins with log2 (fold change **1a/2)** ≥ 1 and p-values <0.05, we established three groups of constrained G4 interacting proteins (Figure 4A). The first and largest group is composed of 204 proteins found significantly enriched on construction **2** (*i.e.* antiparallel topology with two lateral loops) compared to construction **1a** (*i.e.* parallel topology without loop). The second group comprises 190 proteins without significant enrichment on one particular conformation (−2< fold change **1a/2** <2). Finally, we found a third group with only 31 proteins significantly enriched on construction **1a** relatively to structure **2** (Figure 4A and Supplementary Tables 4.1, 4.2 and 4.3). KEGG analysis of the first group and Common proteins showed that both groups define similar significant enriched pathways than those covered by total constrained G4 interacting proteins (Spliceosome, mRNA surveillance, RNA degradation, DNA replication and repair) (Supplementary Figure S4). Surprisingly, the third group of proteins enriched on constrained G4 structure **1a** defines a unique and highly enriched KEGG pathway cluster (p < 0.0005) consisting of factors involved in aminoacyl-tRNA biosynthesis ^50^. Further analysis based on the STRING protein-protein interactions database ^51^ unveiled that all of the components of the multi-tRNA synthetase complex (MSC) are enriched on constrained G4 **1a** (Figure 4B-C). Indeed, in addition to eight cytoplasmic aminoacyl tRNA synthetase enzymes (methionyl MARS, glutaminyl DARS, lysyl KARS, arginyl RARS, isoleucyl IARS, leucyl LARS, aspartyl DARS and glutamyl-, prolyl EPRS-tRNA synthetase) composing this complex, our pull down assay also isolated the three non-enzyme components (AIMP1, AIMP2 and AIMP3, also known as EEF1E, proteins) of the MSC complex (Figure 4B-C). Furthermore, western blotting analysis showed that AIMP1 interacts with constrained and unconstrained G4 structures *in vitro* (Figure 5).

**Figure 4:**
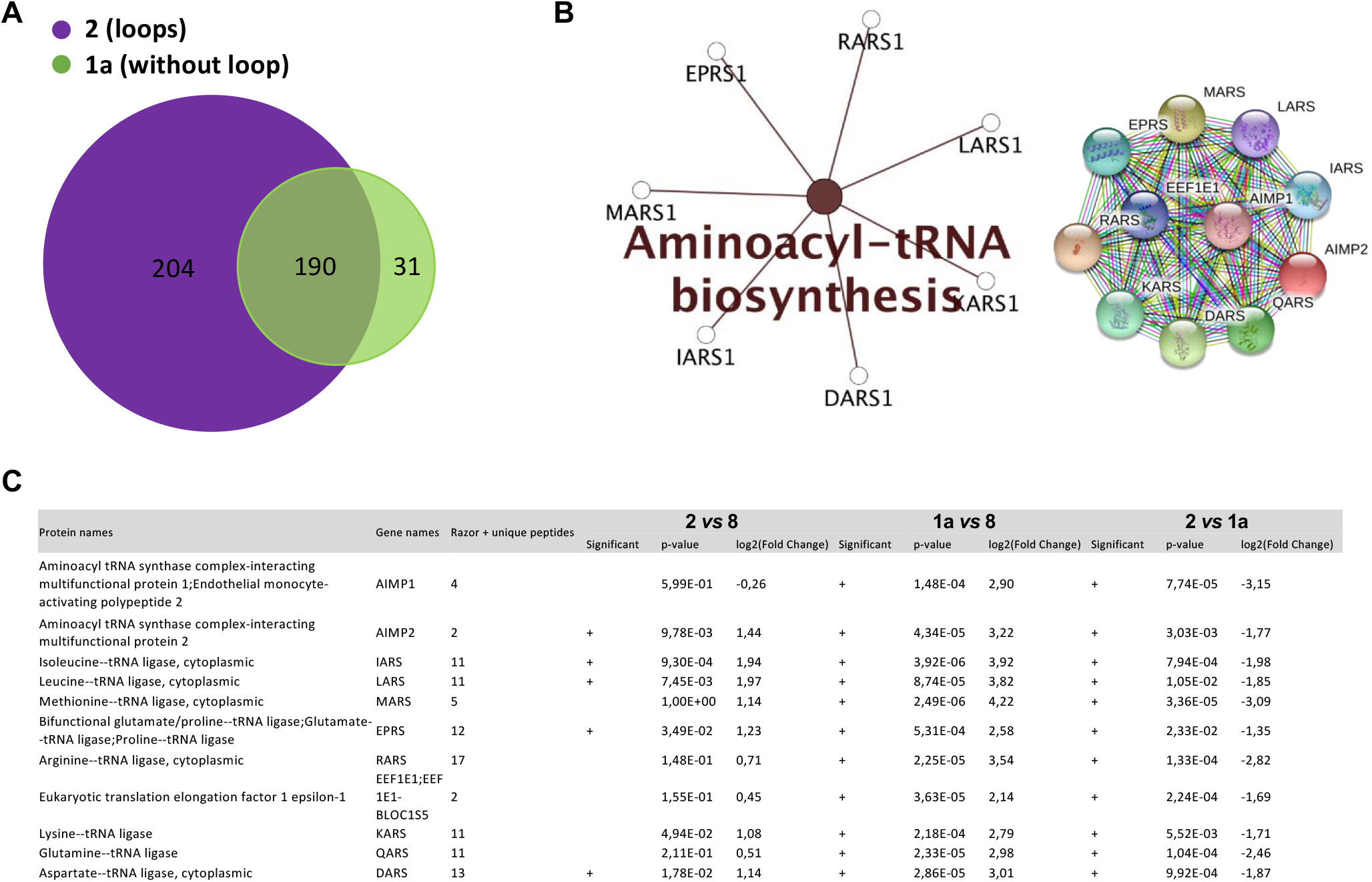
Differential interaction of human proteins with constrained G4 structures adopting different topologies. **A** Diagram showing the differential enrichment of human proteins on **1a** (green) and **2** (purple) constructions. Differential enrichment of proteins on structures **1a** or **2** was determined through statistical analysis using the fold change ≥ 2 and p-value < 0.05, allowing to reach a false discovery rate (FDR) inferior to 5%. **B** KEGG pathway covered by the 32 proteins found enriched on constrained G4 structure **1a** and functional interaction network analysis using STRING ^51^ for the 32 proteins found enriched on construction **1a**. A right-sided (Enrichment) test based on the hyper-geometric distribution was performed on the corresponding Entrez gene IDs for each gene list and the Bonferroni adjustment (p<0.05). **C** MS-based quantitative proteomic analysis of the interaction of MSC-complex proteins with contrained-G4 structures (extracted from Supplementary Table 1). Differentially interacting proteins were sorted out using a fold change ≥ 2 and p-value < 0.05, allowing to reach a false discovery rate (FDR) inferior to 5% according to the Benjamini-Hochberg procedure.

**Figure 5:**
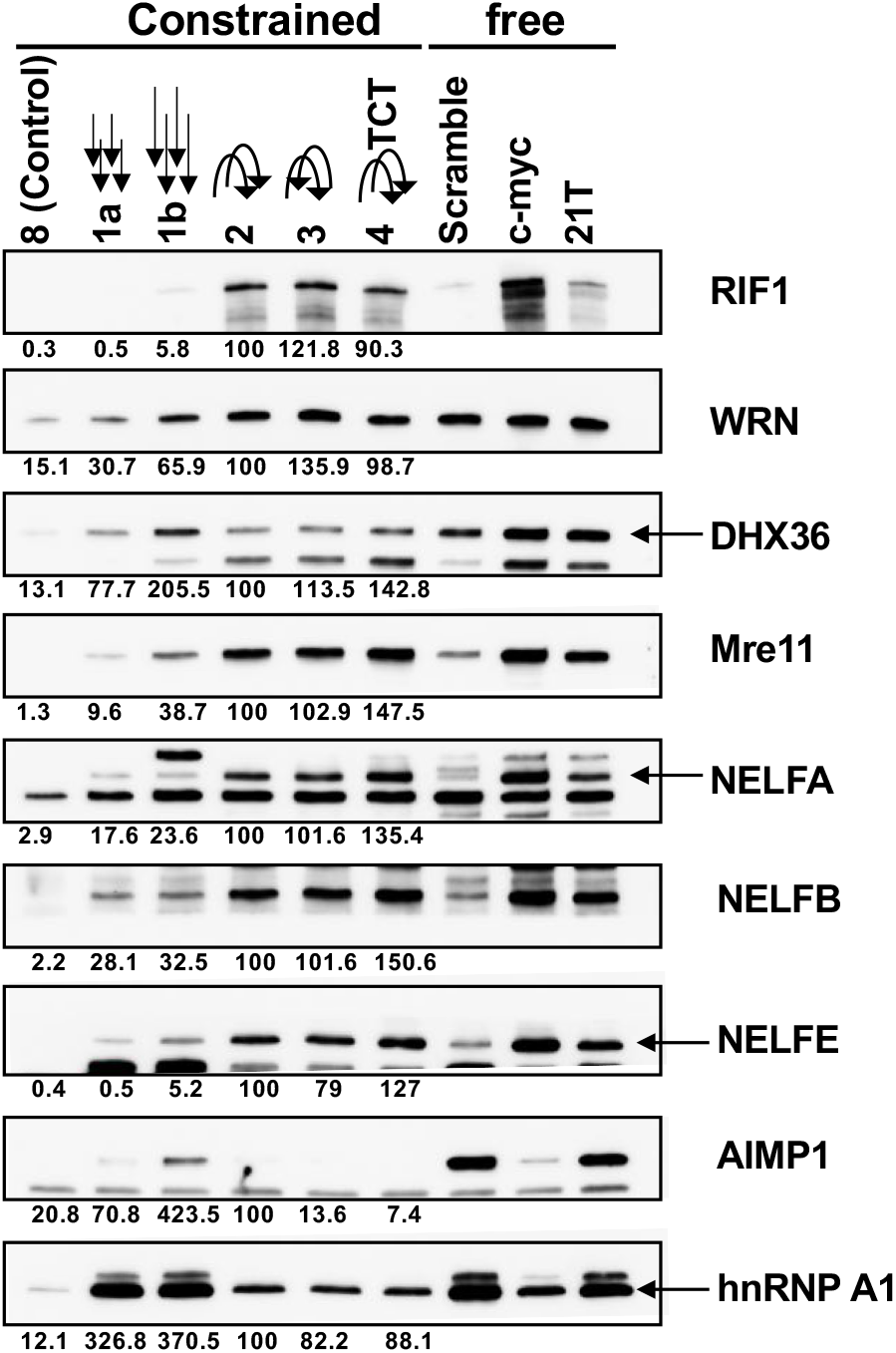
Impact of the orientation and nucleotide composition of connecting loops on the differential enrichment of proteins on G4 structures. Western-blotting analysis and quantification of the interaction of proteins found enriched on constrained G4 structures with modified molecules (**1a, 1b, 2, 3, 4**) and with unconstrained G4 (c-myc and 21T). Arrows indicate the 5’-3’ strand orientation of single-stranded extensions or connecting loops present on different systems. The modification of the nucleotide composition of connecting loops in the system **4** is indicated by the sequence TCT. Construct **8** and scramble sequence were used as control for pull-down performed with constrained of free-G4 structures, respectively.

### Impact of orientation and nucleotide composition of connecting loops on the differential association of proteins on G4 structures

Since most of the proteins characterized in this study are associated with construction **2** (*i.e.* antiparallel with two lateral loops), we explored the impact of the terminal single-stranded extension, loop sequence and loops orientation on the binding of proteins to constrained G4 molecules. First, western blotting analysis of the binding of nine selected factors, found enriched on constrained G4 structures relatively to duplex control **8**, indicates that the relative orientation or the nucleotide loop composition has a not major impact on the binding of these proteins to constrained G4 structures (Figure 5). Indeed, we found that protein signals obtained with systems **2** (*i.e.* with both loops oriented in the same 5’-3’ sense), **3** (*i.e.* with loops oriented in the 5’-3’ and 3’-5’ senses, respectively), and **4** in which the ATT sequence of the external loops was replaced by a TCT, are not significantly different (Figure 5). Next, in order to explore the impact of the extension length, we constructed a new system **1b** with a three-nucleotide extension consisting of the 5’ TTA 3’ sequence. Western-blotting analysis shows that the addition of a supplementary nucleotide on the terminal single-stranded extension of construction **1b** significantly improves the binding of some factors (WRN, DHX36, Mre11, NELF-E and AIMP1). However, intensity signals observed on constrained G4 structures with connecting loops (*i.e.* systems **2-4**) remain considerably stronger relatively to the signals obtained with both **1a** and **1b** constructions, except for AIMP1 (Figure 5). These results indicate that longer single-stranded extensions improve the binding of proteins to type **1** constructions. Unconstrained G4 structures (free) formed by the telomeric and the c-myc promoter sequences were also used to further characterize their interaction (Figure 5, right part). Most of the factors enriched on constrained G4 structures also present a significant interaction with non-constrained G4-forming sequences, relative to the scramble sequence, indicating a selective binding of these proteins to G4 structures. However, proteins showing a significant interaction with **1a** and **1b** systems (hnRNP A1, AIMP1, WRN and at lesser extent DHX36) also display a strong interaction with the scramble non-G4 forming oligonucleotide, confirming their ability to bind both G4 structures and single-stranded unfolded G-rich DNA.

### Constrained G4 structures identify the NELF complex as a new G4 interacting factor

MS-based quantitative proteomic analyses identified NELF-A, NELF-B, NELF-E and NELF-C/D, all members of the NELF complex ^52^, as enriched on constrained G4 structures relatively to the duplex control construction **8** (Figure 6A, supplementary Table 2). Selective interaction of NELF complex proteins with G4 structures was further confirmed through western blotting analysis performed using both constrained and unconstrained G4 structures (Figure 5). In order to further characterize the interaction of the NELF complex with G4 structures, we performed the reverse experiment in which an immunoprecipitated NELF complex was used to investigate its interaction with constrained G4 structures. For that, the NELF complex was immunoprecipitated from a HeLa cell line overexpressing an ectopic Flag-tagged form of the NELFE protein. After elution by competition with a Flag peptide (Supplementary Figure S5), the NELF complex was use in pull-down experiments using constrained constructions and western blotting analyses were performed to quantify the relative signal of the bound NELFE protein. As shown in the Figure 6B, the NELFE protein is significantly enriched on constrained G4 structures relatively to the cyclopeptide (CP-T23) and duplex control **8**, indicating a selective binding of the NELFE protein to G4 structures. In addition, hybridization with an anti-NELFA antibody indicates that at least two proteins from the immunoprecipitated NELF complex are found enriched on constrained G4 structures relatively to the controls (CP-T23 and system **8**) (data not shown).

**Figure 6:**
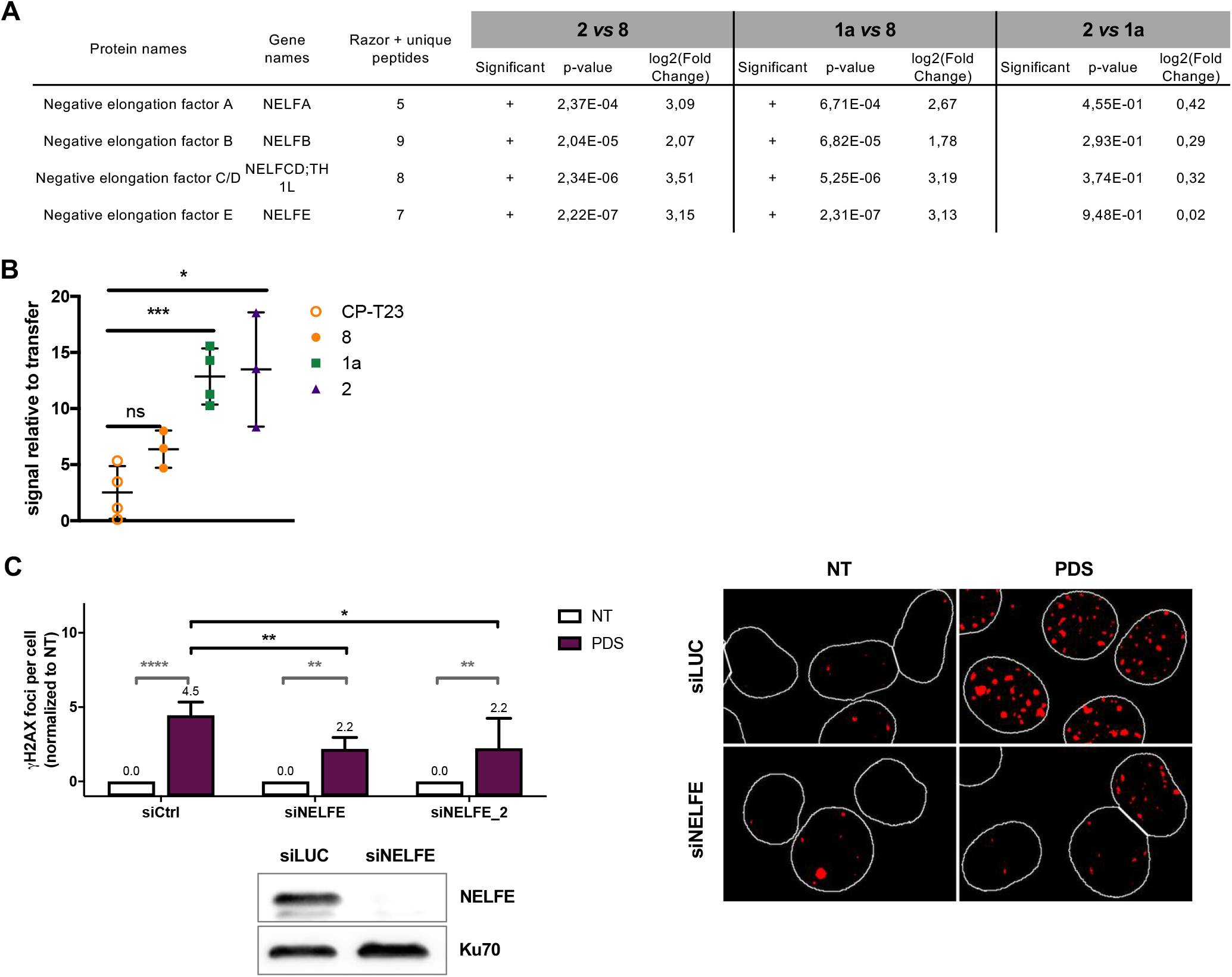
NELF complex interact with G4 structures and modulates the cellular response to G4 ligands. **A** MS-based quantitative proteomic analysis of the interaction of the NELF-complex proteins with contrained-G4 structures (extracted from Supplementary Table 1). Differentially interacting proteins were sorted out using a fold change ≥ 2 and p-value < 0.05, allowing to reach a false discovery rate (FDR) inferior to 5% according to the Benjamini-Hochberg procedure. **B** Quantification of the interaction of immunoprecipitated Flag NELF-E protein with constrained G4 structures (**1a, 2**) relative to cyclopeptide (CP-T23) and duplex control (**8**) constructions. Error bars represent SD from the means, *n* ≥ 3 independent experiments. *p* values were calculated using unpaired t-tests (without corrections for multiple comparisons). ns: p > 0.05; *: p < 0.05; **: p < 0.01; ***: p < 0.001; ****: p < 0.0001. ns non-significant difference. **C** Quantification and representative images gH2AX foci fluorescence signal (red) detected HeLa cells transfected with control (Ctrl), or two NELF-E siRNAs (independent sequences) and treated with PDS (20 μM) for 4 h. Error bars represent SD from the means, *n* ≥ 3 independent experiments. *P* values were calculated using an unpaired multiple Student’s *t* test. ns: p>0.05; *: p<0.05; **: p<0.01; ***: p<0.001; ****: p<0.0001signals. Western-blotting analysis of NELF-E depletion in HeLa cells following siRNA treatment is shown.

### The NELF complex facilitates DNA double-strand breaks induction by Pyridostatin

In human cells the NELF complex plays an essential role in the RNA-Pol II pausing mechanism ^53, 54^. RNA-Pol II pausing is a highly controlled mechanism regulating gene expression in eukaryotic cells ^53^. During the pausing state, RNA-Pol II remains tightly associated with nascent RNA molecule in a promoter proximal region. Bioinformatic and pangenomic studies clearly establish that the formation of G4 structures, which are significantly enriched in promoter regions in human cells, correlates with the formation of R-loops and RNA-Pol II pausing sites, two processes associated with RNA-Pol II arrests and the induction of transcription dependent double-stranded DNA breaks (DSBs) ^30-32, 55-60^. Pyridostatin (PDS), one of the most selective and potent G4 stabilizers described so far, provokes a rapid induction of transcription dependent DSBs in human cells (^61, 62^ and unpublished data). In order to investigate the impact of the NELF complex on PDS-induced DSBs, we quantified γH2AX signals (DSBs marker) through immunofluorescence studies in NELF proficient and deficient (siRNA-depleted NELF-E) HeLa cells. As shown in figure 6C, depletion of the NELFE protein, which also leads to the reduction of other NELF proteins in human cells ^52, 63^, provokes a significant reduction of PDS-induced DSBs signals relatively to the control cells. This result indicates that the NELF complex and RNA Pol II pausing favours DSB induction following G4 stabilization.

## Discussion

The relationship between G4 structures/motifs and the progressive discovery of proteins that modulate the dynamic of G4 formation, such as helicases or other proteins involved in DNA and RNA transactions has expanded our knowledge to understand the ubiquitous function of G4 structures on cellular metabolism and cell fate ^5, 6, 13, 15, 17-20, 64^. In the present study, we identified through a MS-based quantitative proteomic analysis 425 human proteins significantly enriched on constrained G4 structures relative to a duplex DNA control, in which well-described G4 interacting factors were present. However, we found that some of well-known G4-interacting proteins, such as BLM and WRN, are not enriched on constrained G4 relatively to duplex control, thereby confirming the previously reported non-selective interaction of these proteins with G4s relative to duplex or hairpin DNAs ^65^. Interestingly, numerous proteins interacting with constrained G4-DNA structures identified in this work are involved in RNA transactions: splicing, RNA degradation, transport and RNA surveillance pathways. Moreover, about one hundred proteins enriched from 425 total proteins identified here on constrained G4-DNA structures have been already identified in similar works using RNA G4 structures to trap G4-RNAs binding factors ^35–37^. It is noteworthy that most RNA Binding Proteins (RBP) interact with RNAs through OB-fold, RGG, RRM and GARD motifs that have been also involved in the recognition of G4 structures ^21, 22, 66^. Most G4 interacting proteins unfold G4 in order to counteract their impact on DNA and RNA transactions. Interestingly, 62 out of the 425 proteins identified through our approach have also been characterized as playing an important role in the resistance to cytotoxicity induced by G4-ligands ^49^, and for many of them our work provides an evidence for a direct or indirect interaction with G4.

An important finding from our results concerns the differential enrichment of G4 interacting proteins on constrained G4 molecules mimicking parallel (systems **1, 6** and **7**) *versus* antiparallel conformations (systems **2-5**). MS-based quantitative proteomic analysis identified 204 proteins significantly enriched on the G4 construction **2** relative to the G4 construction **1**, and only 31 proteins with an inverse selectivity. Surprisingly, relative strand orientation and nucleotide composition of terminal connecting loops had no major impact on the binding of proteins to loops containing constructions (**2-4**), suggesting either a minor role of loops for the interaction with proteins recognizing antiparallel constrained G4, or a role related to their folding into a particular conformation independent on nucleotide composition or strand orientation. In agreement with the first hypothesis, selective binding of proteins and small molecules to G4 structures are driven mainly through π-π interactions with tetrad faces, although additional contacts with lateral grooves and connecting loops have been shown to stabilize both molecules and protein-G4 complexes ^22, 67^. Also consistent with the first hypothesis, proteins found enriched on system **2-4** relatively to system **1a-b**, bind unconstrained G4 structures formed by the c-myc and telomeric sequences. In solution, both sequences have been shown to adopt different topologies and exhibit loops with different nucleotide compositions and strand orientations ^68, 69^. Altogether, these results indicate that the interaction with tetrad faces would be a major determinant for the binding of human proteins to system **2-4**. Furthermore, no significant difference was found for 190 out of the 425 proteins interacting with systems **1a** and **2**, a result that would indicate that these proteins interact with a common structural motif. Finally, although the addition of a supplementary nucleotide to construction **1a** significantly increases the binding of system **2** enriched proteins on the system **1,** we assume that a part of the differential binding of human proteins to loops-containing systems **2-4** relative to loop-free constructions **1, 6** and **7** would be dependent on the accessibility of the tetrad face or on supplementary contacts with lateral grooves. Future studies with constrained structures containing propeller loops will be needed to fully characterize this impact.

An intriguing finding from our study is the characterization of the multi-tRNA synthetase complex (MSC) as a G4-interacting complex. The interaction of the MSC complex with G4 structures was confirmed by a recent publication that identified the MSC proteins in a molecular screen identifying RNA G4-interacting factors ^36^. The MSC complex consists of eight Aminoacyl-tRNA Synthetases (ARSs) and three non-enzymatic ARS-interacting multi-functional proteins (AIMP1/p43, AIMP2/p38, and AIMP3/p18) that play an essential role in the protein synthesis by catalyzing the activation of amino acids and linking them to their cognate transfer RNAs (tRNAs) ^50^. AIMP1/p43 protein is a multifunction protein involved in various physiological and pathological processes. AIMP1 is the precursor of EMAP II, which was released after AIMP1 cleavage ^70^. Surprisingly, the C-terminal segment of EMAP II are 50% identical to residues 205-364 of p42 protein from *S. cerevisiae* that was firstly identify as G4p1 a protein that shows a high and specific affinity for G4 nucleic acids ^71^.

Finally, the interaction of NELF complex with G4 structures is a major finding of our study. To the best of our knowledge, this is the first time that a physical interaction between the NELF complex and G4 is reported. The NELF complex comprises the NELFA, NELFB, NELFC/D and NELFE proteins. Whether the interaction of NELF with G4 structures is direct or indirect needs to be elucidated. Nevertheless, the NELFE protein contains a RRM-domain (RNA Recognition Motif) ^52^ that is present in different RNA binding proteins and interacts with G-rich tracts. These RRM motifs have also been involved in the interaction of several proteins with G4 structures, such as hnRNP A1, and in agreement with our findings, a NELF sub-complex formed by NELFA and NELFE proteins has been shown to selectively bind GC rich sequences ^72^. Numerous bioinformatic and genomics analysis have established a strong correlation between G4 and RNA Pol II pausing occurring at the promoter proximal regions ^31, 73^. Transcriptional pausing consists in the arrest of RNA-Pol II mainly in a promoter proximal position and acts as a transcriptional checkpoint regulating gene expression ^53^. It is induced by the association of two complexes, NELF and DSIF, with RNA-Pol II at the early stages of transcriptional elongation ^54^. Although G4 motifs are found at pausing sites ^31^, it is still unclear whether G4s act as signals for RNA-Pol II pausing. However, during transcription the impact of G4 on RNA Pol II progression could reflect their capacity to act as physical barriers, to promote the formation of other secondary structures such as R-loops or finally to drive the binding of protein complexes regulating RNA Pol II progression. In human cells, G4 motifs and RNA Pol II pausing have been also associated with transcription-dependent DNA breaks ^31, 32, 55^, and the stabilisation of G4 structures by small molecules has been shown to provoke the formation of double-stranded DNA breaks that are, in part, dependent on RNA-Pol II transcription ^61^. In this study, we show that the NELF complex regulates the formation of double-stranded DNA breaks induced by pyridostatin, a potent and selective G4 ligand. Altogether, our results suggest that G4 formation could facilitate the binding of the NELF complex to the chromatin to promote RNA Pol II pausing, and thus to act as a mediator of the response to G4 stabilization in human cells.

In this study, we show that biotin-functionalized constrained nucleic acids structures are powerful tools to identify proteins interacting with non-canonical secondary structures such as G4. We validated our approach by the identification of well-known G4 interacting factors and the functional characterization of new protein complexes related to G4 metabolism. Especially, the identification of NELF complex as a G4 interacting factor establishes a physical link between G4 structures and RNA Pol II pausing mechanism.

Future directions of our approach will concern the construction of constrained G4 structures mimicking intramolecular parallel G4 structures in order to refine the impact of propeller loops on protein binding. Finally, other constrained nucleic acid constructions may represent powerful tools to identify proteins interacting with other non-canonical secondary structures such as the i-motif or R-loops.

## Material and Methods

### Constained G4 and HP synthesis

The different constrained systems **1-8** were prepared according to the previously reported protocols (57-60). Biotinylated c-myc (5’GGA-GGG-TGG-GGA-GGG-TGG-GGA-A-TEG-biot), 21T (5’ TTA-GGG-TTA-GGG-TTA-GGG-TTA-GGG-TT-TEG-biot) and scramble (5’ AAG-TGT-GTG-TGT-GTG-TGT-GTG-TGA-AG-TEG-biot) sequences were purchased from Eurogentec.

### Cell culture

HeLa cells were grown in humidified atmosphere with 5% CO2 at 37°C, in Dulbecco’s Modified Eagle Medium (Gibco). Culture medium was supplemented with 10% fœtal bovine serum (Eurobio), 100 U/mL penicillin (Gibco) and 100 μg/mL streptomycin (Gibco).

### Plasmid constructions Cell transfection and transduction

The pCMVsVg (envelope plasmid) were kindly provided by E. Gilson. pLPC-puro-N-Flag was a gift from Titia de Lange (Addgene plasmid # 12521; http://n2t.net/addgene:12521; RRID: Addgene_12521). The pLPC-puro-N-Flag-NELFE plasmid was obtained by amplification of the human NELFE cDNA (obtained from gene synthesis, GeneArt, LifeTechnlogies) using oligonucleotides: Fwd 5’-GACGATGACGATAAAGGATCCTTGGTGATACCCCCCGGACT-3’ and Rev-5’CCCTCTAGATGCATGCTCGAGCTAGAAGCCATCCACAAGGTTTTCC-3’ and inserted in the pLPC-puro-N-Flag plasmid between XhoI and BamH1 restriction sites. Retroviral production was performed by transient transfection of HEK-GP2 293 cells (Clontech) with 0.8 μg pCMVsVg and 1.2 μg of pLPC-puro-N-Flag-NELFE plasmids using JetPrime reagent (polyplus). Retroviral particles were recovered from culture supernatant of HEK-GP2 293 cells 48 h and 72 h post transfection. For transduction, fifty thousand Hela cells were plated on six-well plates 24 h prior to transduction. Cell population was selected by puromycin resistance (1 μg/ml). The expression of Flag-NELF-E protein was verified by western blotting with Flag and NELF-E antibodies.

### Pull-down from total protein extract with constrained G4s

First, proteins were bound to constrained G4s. Briefly, 1 mg of NHEJ protein extract (prepared as previously described ^46^) was incubated with 10 μM of one of constrained G4s (Figure A) in 100 μL of binding buffer (20 mM Hepes (pH 7.5), 50 mM KCl, 0.01% NP40 and 0.5 mM EDTA) for 1.5 h at 4 °C under intermittent shaking (10 sec at 1400 rpm every 2 min). During protein-G4 binding step, 1 mL of streptavidin-coupled magnetic beads (Promega, Z5481) per condition were washed three times during 10 min at 4°C (under intermittent shaking) with binding buffer. Then, 100 μL of protein-G4 binding solution was put onto washed streptavidin-coupled magnetic beads for 30 min at 4°C under intermittent shaking. After this step, supernatant was stored at −80°C as “unbound fraction” and beads were washed during 10 min three times at 4 °C (under intermittent shaking) with 0.1% NP40, 150 mM NaCl PBS. Finally, beads were incubated with 100 μL of 0.01% bromophenol blue, 15% glycerol, 2% SDS, 60 mM Tris-HCl (pH 8) for 10 min at 95 °C and supernatant were collected and stored at −80°C before being used in mass spectrometry and western blotting assays.

### MS-based quantitative proteomic analysis

Eluted proteins were stacked in a single band in the top of a SDS-PAGE gel (4-12% NuPAGE, Life Technologies) and stained with Coomassie blue R-250 before in-gel digestion using modified trypsin (Promega, sequencing grade) as previously described ^74^. Resulting peptides were analyzed by online nanoliquid chromatography coupled to tandem MS (UltiMate 3000 and LTQ-Orbitrap Velos Pro, Thermo Scientific). Peptides were sampled on a 300 μm × 5 mm PepMap C18 precolumn and separated on a 75 μm × 250 mm C18 column (PepMap, Thermo Scientific) using a 120-min gradient. MS and MS/MS data were acquired using Xcalibur (Thermo Scientific).

Peptides and proteins were identified and quantified using MaxQuant (version 1.6.2.10, ^75^) using the Uniprot database (*Homo sapiens* reference proteome, October 22^nd^ 2018 version) and the frequently observed contaminant database embedded in MaxQuant. Trypsin was chosen as the enzyme and 2 missed cleavages were allowed. Peptide modifications allowed during the search were: carbamidomethylation (C, fixed), acetyl (Protein N-ter, variable) and oxidation (M, variable). Minimum peptide length was set to 7 amino acids. Minimum number of peptides and razor + unique peptides were set to 1. Maximum false discovery rates - calculated by employing a reverse database strategy - were set to 0.01 at peptide and protein levels. The mass spectrometry proteomics data have been deposited to the ProteomeXchange Consortium via the PRIDE ^76^ partner repository with the dataset identifier PXD 021003.

Statistical analysis were performed using ProStaR ^77^. Proteins identified in the reverse and contaminant databases, proteins only identified by site, proteins identified with only 1 peptide and proteins exhibiting less than 3 intensity values in one condition were discarded from the list. After log2 transformation, intensity values were normalized by median centering before missing value imputation (slsa algorithm for partially observed values in the condition and DetQuantile algorithm set to first percentile for totally absent values in the condition); statistical testing was conducted using limma test. Differentially interacting proteins were sorted out using a log2 (fold change) cut-off of 1 and a p-value cut-off allowing to reach an FDR inferior to 5% according to the Benjamini-Hochberg procedure.

### Immunoprecipitation of Flag-NELF-E and pull-down with constrained G4s

Four days after seeding, around 30.10^6^ Flag-NELF-E expressing HeLa cells were harvested by scrapping in cold PBS. After centrifugation at 2000 g for 5 min at 4 °C, pelleted cells were lysed for 40 min at 4 °C with 1 mL of non-denaturing lysis buffer (50 mM Hepes (pH 7.5), 150 mM NaCl, 0.01% NP40) complemented with 1 mM DTT, 1X Halt Protease Inhibitor Cocktail (ThermoScientific), 23 U/mL benzonase. Cells were then centrifugated at 12000 rpm at 4 °C for 10 min and protein concentrations in total extract were determined by measuring absorbance at 280 nm (Nanodrop). From total extract, 1 mg of proteins was used to performed incubation overnight at 4°C with 5 μg of anti-Flag mouse antibody (Sigma) in 250 μL of complemented non-denaturing lysis buffer. Protein A/G-coupled magnetic beads (Pierce, 88802) were washed with complemented non-denaturing lysis buffer and 25 μL of beads were used to immunoprecipitated Flag antibody for 1h at 4 °C. Supernatant was collected as “flag unbound fraction” and protein concentration was determined as described. Immunoprecipitated proteins were eluted overnight at 4 °C with 50 μg of Flag peptide (Sigma-Aldrich) in 100 μL complemented non-denaturing lysis buffer. Supernatant was collected as “flag eluted fraction” and protein concentration was determined as described. For protein-G4 binding step, 7 μg of flag eluted fraction were incubated with 1 μM of constrained G4s in 100 μL of binding buffer overnight at 4 °C, and pull-down was performed as described above. Finally, beads were incubated with 30 μL of 0.01% bromophenol blue, 15% glycerol, 2% SDS, 60 mM Tris (pH 8) for 10 min at 95 °C and supernatant were collected and stored at −80°C before being used for western blotting assays.

### Western Blotting

For total pulled-down proteins and for Flag-eluted pulled-down proteins, respectively 45 μL and 10 μL of proteins were loaded for each condition and separated on gradient 4-12% polyacrylamide TGX Stain-Free pre-cast gels (Biorad) and transferred onto nitrocellulose membrane (Biorad). Before blocking (0.1% Tween-20, non-fat dry milk 5% and PBS), UV exposition of membrane was used to confirm homogeneous loading and to quantify transfer signal. The membrane was successively probed with primary antibodies and appropriate goat secondary antibodies coupled to horseradish peroxidase (described in table below). Chemidoc imager (Biorad) was used to perform UV and Clarity ECL (Biorad) detection. Digital data were processed and quantified using ImageLab (Biorad) or ImageJ softwares. Quantifications of antibody signal are relative to transfer signal.

| Target | Dilution | Species | Class | Reference | Manufacturer |
| --- | --- | --- | --- | --- | --- |
| KU70 | 0.2 μg/mL | Mouse | Monoclonal | MA5-13110 | Invitrogen |
| eIF4G | 1/1000 | Rabbit | Polyclonal | 2498 | Cell Signaling Technology |
| WRN | 1/2000 | Mouse | Monoclonal | W0393 | Sigma-Aldrich |
| Mre11 | 1/1000 | Mouse | Monoclonal | 611366 | BD bioscience |
| hnRNP A1 | 1/1000 | Rabbit | Polyclonal | GTX106208 | Gene Tex |
| DHX36 | 1/1000 | Rabbit | Polyclonal | HPA035399 | Sigma-Aldrich |
| CNBP | 8/1000 |  |  |  | Kindly provided by N. Calcaterra's group |
| NCL23 | 1/1000 | Rabbit | Polyclonal | ab50279 | Abcam |
| NELFE | 1 μg/mL | Rabbit | Monoclonal | A301-913A | Bethyl laboratories |
| NELFB (COBRA) | 1 μg/mL | Rabbit | Monoclonal | A301-911A | Bethyl laboratories |
| Anti-rabbit | 1:10000 | Goat | Polyclonal | 111-035-003 | Jackson Immunoresearch |
| Anti-mouse | 1:10000 | Goat | Polyclonal | 115-035-003 | Jackson Immunoresearch |

### RNA interferences

HeLa I3 cells were seeded at 250.000 cells per well in a 6-wells plate. siRNA (Table) were transfected twice at 50 nM final concentration per well with Lipofectamine RNAiMax Reagent (Invitrogen) according to manufacturer’s instructions. Cells were treated and proteic extracts are realized 72 h after the first transfection.

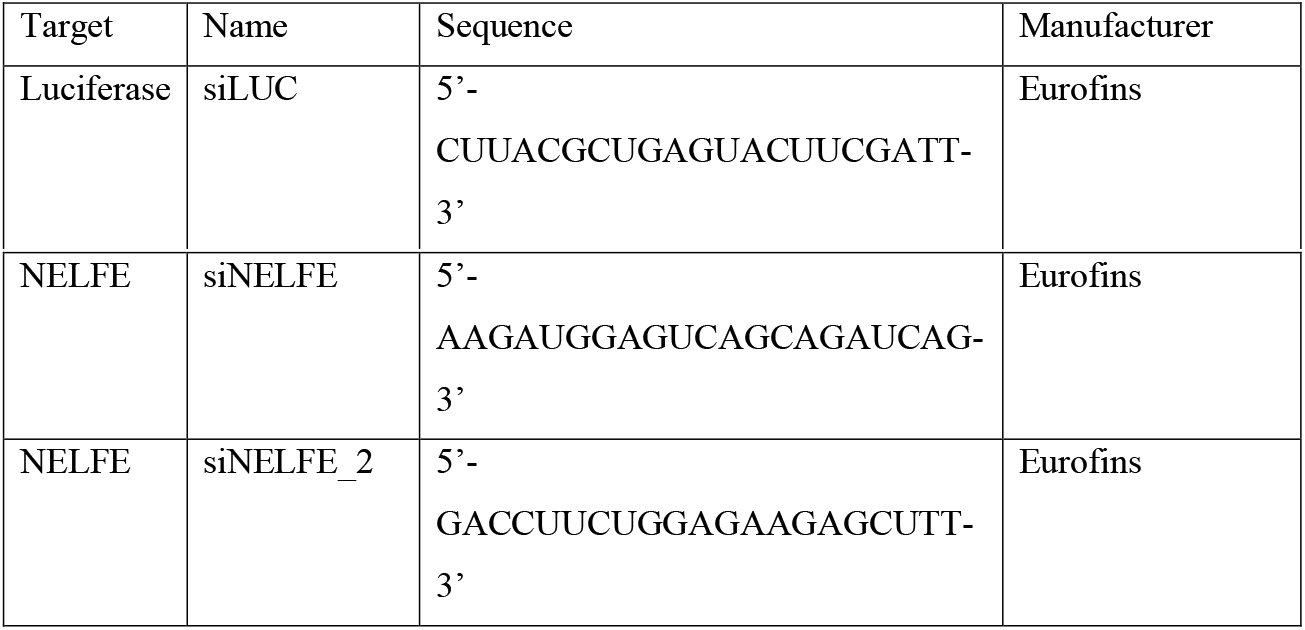

### Immunofluorescence

HeLa cells were seeded in 24-wells plate at 100.000 cells/well on glass coverslips (VWR, #631-0150). Twenty-four hours later, HeLa cells were treated with 20 μM pyridostatin (Sigma-Aldrich; CAS number 1085412-37-8) for 4 h, and then washed with PBS and fixed with 2% paraformaldehyde in PBS at room temperature for 10 min, washed with PBS and permeabilized for 15 min at room temperature with 10 mM Tris-HCl pH 7.5, 120 mM KCl, 20 mM NaCl, 0.1% Triton-X 100. Then, cells were washed with PBS and incubated for about 1 h at 37°C in blocking buffer (20 mM Tris-HCl pH 7.5, 150 mM NaCl, 2% BSA, 0.2% fish gelatin, 0.1% Triton-X 100) prior to incubation overnight at 4 °C with gH2AX (Phospho S139) antibody (Abcam, #81299) diluted at 0.7 μg/mL in blocking buffer. Cells were then washed with 0.1% Tween20-PBS and incubated with secondary goat anti-rabbit antibody coupled to AlexaFluor 488 (Thermo Fisher Scientific) diluted at 2 μg/mL in blocking buffer for 1 h at room temperature. At last, cells were washed with 0.1% Tween20-PBS and stained with 0.1 μg/mL DAPI for 20 min at room temperature, and coverslips were mounted with Vectashield mounting medium (Vector Laboratories). Nuclear gH2AX foci staining overlapping with DAPI staining were quantified with ImageJ software. Quantifications of nuclear gH2AX foci induced by pyridostatin are represented normalized to non-treated (NT) conditions.

### KEGG pathway, gene ontology

The ClueGO v2.3.3 ^78, 79^ plugin for Cytoscape ^80^ (v3.8) was used to determine networks of enriched KEGG pathways and Gene Ontology terms (Biological Process and Molecular Function). A right-sided (Enrichment) test based on the hyper-geometric distribution was performed on the corresponding Entrez gene IDs for each gene list and the Bonferroni adjustment (p < 0.05) was applied to correct for multiple hypothesis testing. The Kappa-statistics score threshold was set to 0.4 and GO term fusion was used to diminish redundancy of terms shared by similar proteins. Other parameters include: GO level intervals (3-8 genes) and Group Merge (50%).

### Statistical analysis

All results provide from at least three independent experiments. Statistical analyses were performed with GraphPad Prism Software (version 8). For gH2AX quantifications analyses, multiple unpaired t-tests (without corrections for multiple comparisons) were performed between pairs of conditions. On all figures, significant differences between specified pairs of conditions are shown by asterisks (*: p-value < 0.05; **: p-value < 0.01; ***: p-value < 0.0005; ****: p-value < 0.0001). NS means non-significant difference.

## Acknowledgments

This work was funded by grants from ANR (ANR-16-CE11-0006-01), and La Ligue Nationale Contre le Cancer (Equipe Labellisée 2018). Patrick Calsou is a researcher from INSERM. Proteomic experiments were partly supported by the ProFi grant (ANR-10-INBS-08-01). The Nanobio-ICMG platform (FR2607) is acknowledged for providing facilities for the synthesis and purification of oligonucleotides (R. Lartia) as well as for mass spectrometry analyses (A. Durand, L. Fort and R. Guéret).

## Supplementary Figures

**Supplementary Table1** MS-based proteomic analyses of proteins interacting with constrained structures **1a, 2** and the control **8** (duplex).

**Supplementary Table2** List of 425 proteins enriched on constrained G4 structures **1a-2** compared to duplex control **8** (fold change ≥ 2 and p-value < 0.05, allowing to reach a false discovery rate (FDR) inferior to 5%).

**Supplementary Table3** List of 214 out of 425 proteins enriched on G4 structures that have been described as nucleic acid interacting factors, as indicated by terms from GO Molecular functions data analyses.

**Supplementary Table4.1** List of 31 proteins found enriched on system 1a relatively to system 2

**Supplementary Table4.2** List of 204 proteins found enriched on system 2 relatively to system 1a

**Supplementary Table4.3** List of 190 proteins Common to system 1a and 2

**Supplementary Figure S1.**
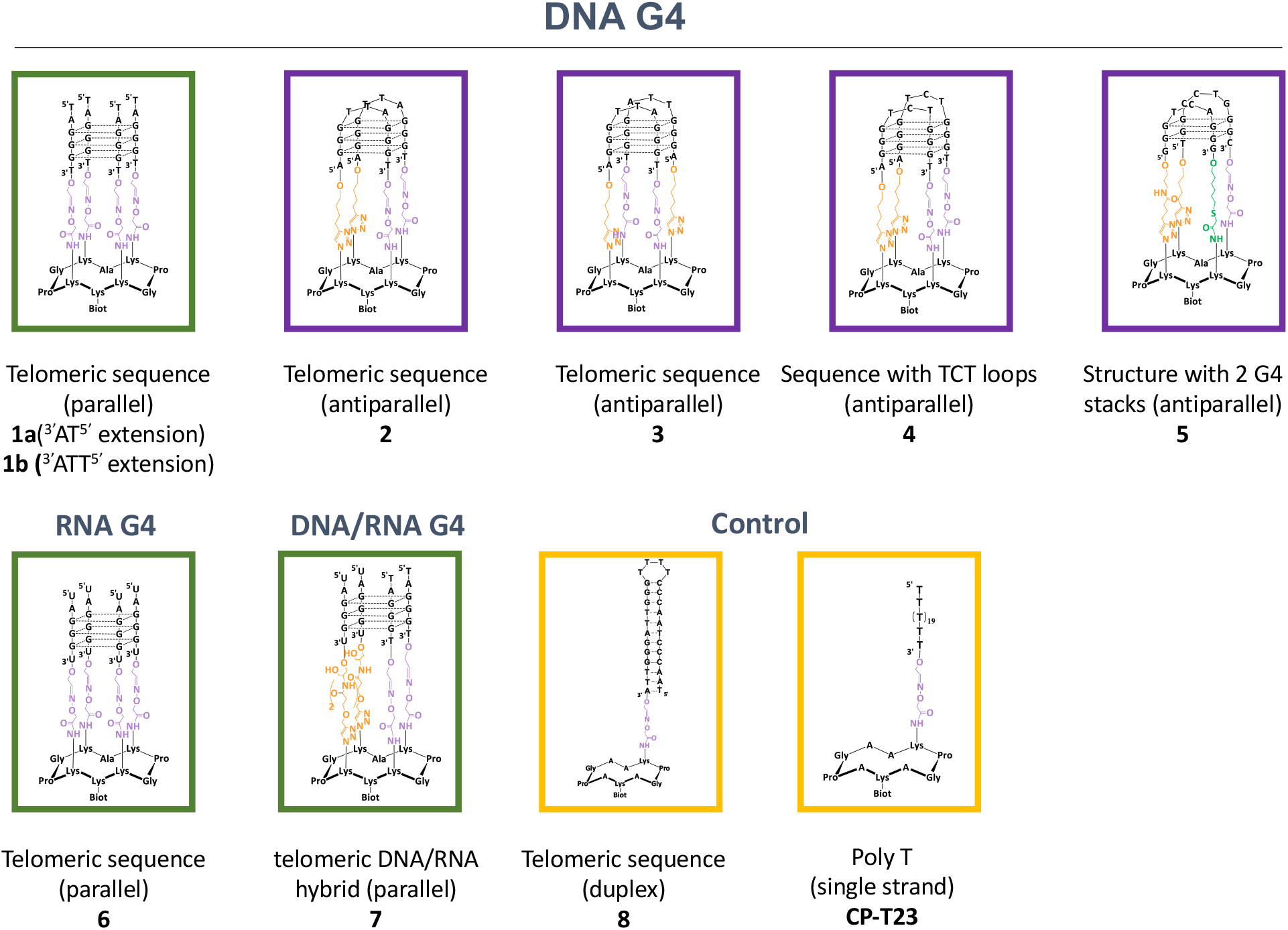
Schematic representation of the constrained molecules used in this study

**Supplementary Figure S2.**
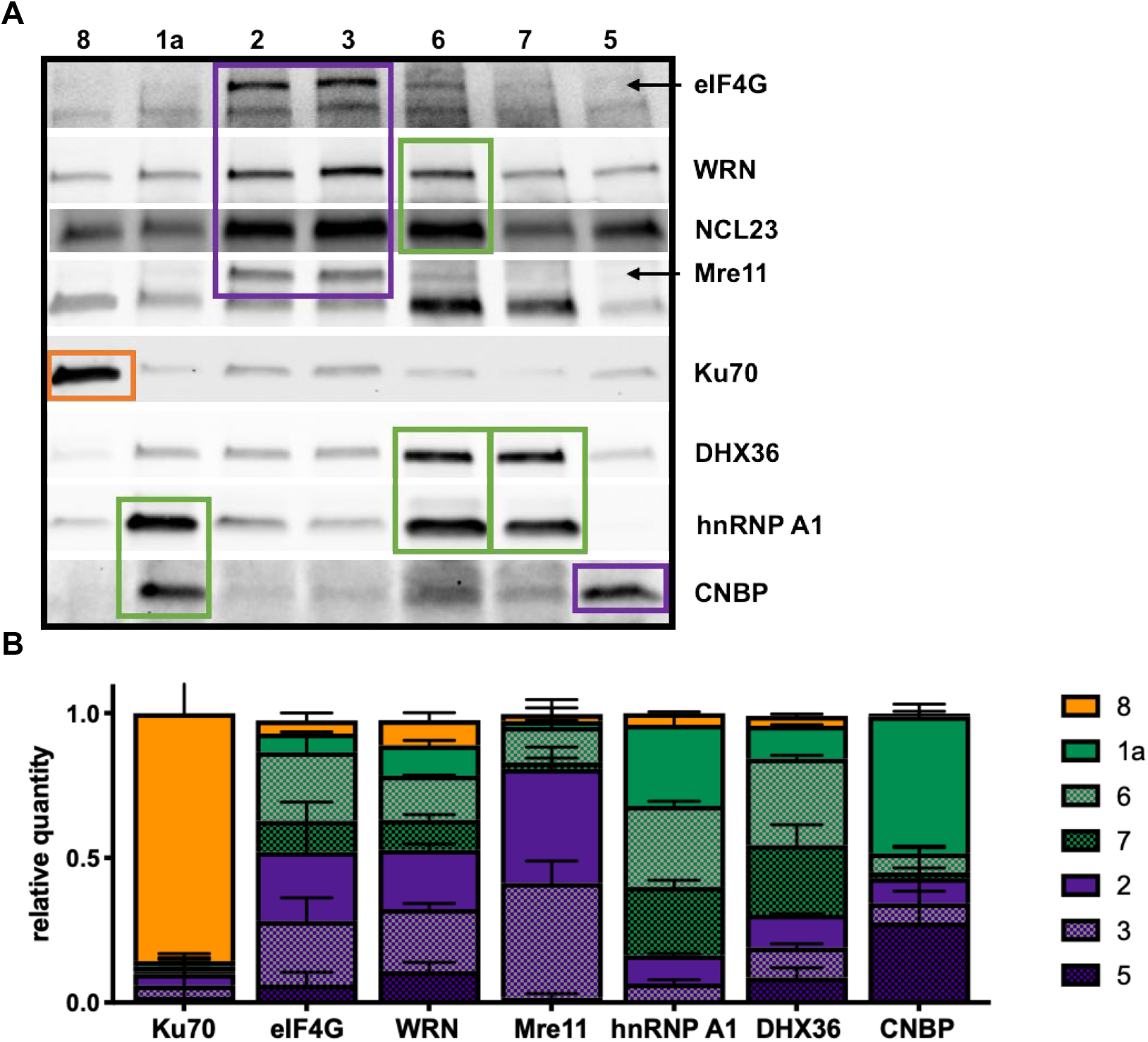
Western-blotting analysis and quantification of molecular interactions of known-G4 interacting factors with constrained G4 structures adopting different topologies (see Supplementary Figure S1). The relative quantity was obtained from the ratio between the signal observed from a particular construction and the sum of signals from all wells. Digital data were processed and quantified using ImageLab (Biorad) or ImageJ softwares. Green, purple and oranges boxes indicate interaction with Parallel **1a**, antiparallel **2** or duplex DNA **8** respectively.

**Supplementary Figure S3.**
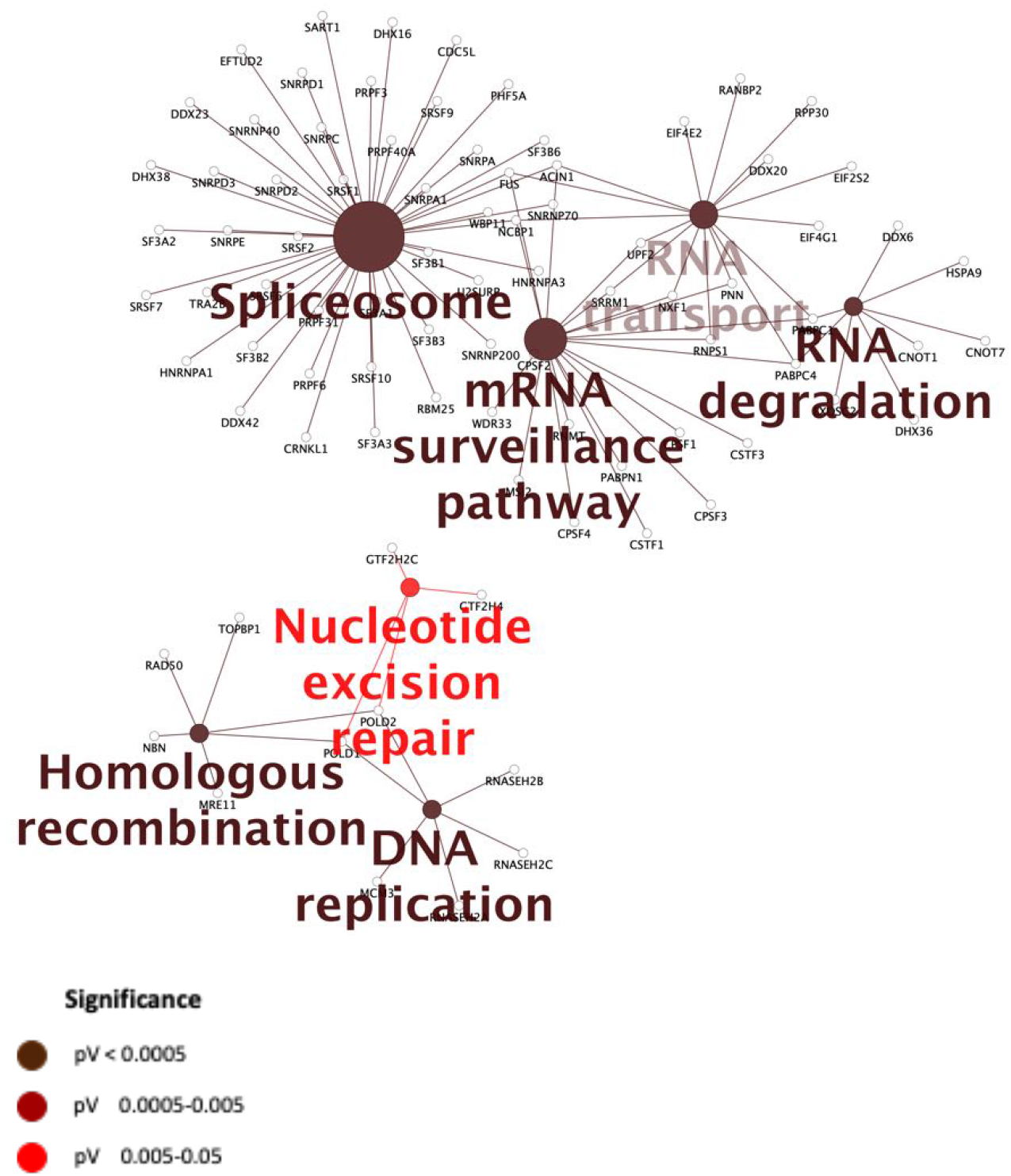
Enriched KEGG pathways for the 214 nuclear proteins interacting with nucleic acids found enriched on constrained G4 structures **1a** and **2**.

**Supplementary Figure S4.**
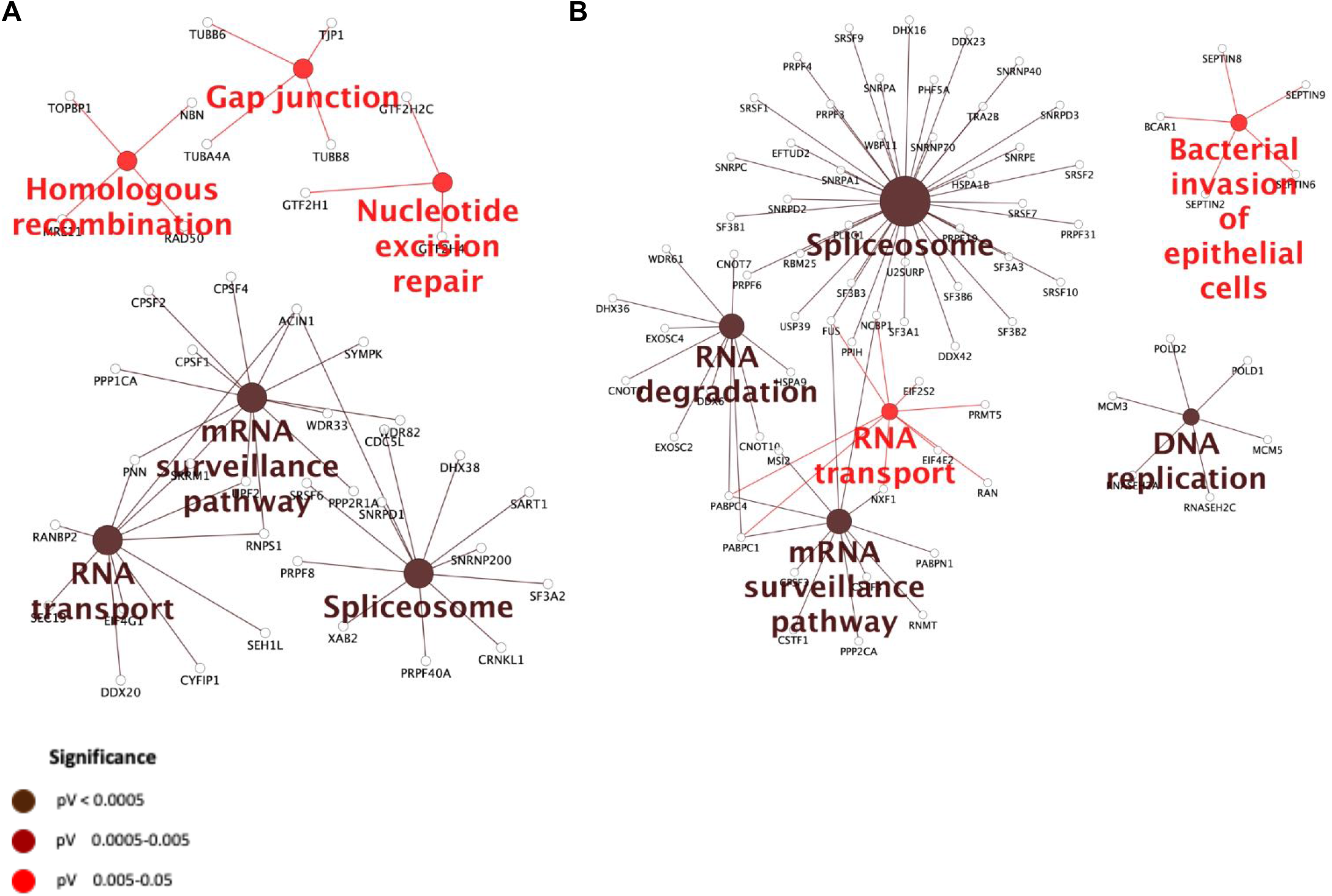
Most significant pathways and processes covered by constrained G4 interacting proteins. Enriched KEGG pathways of (**A**) the 190 proteins present in the Common group and (**B**) the 203 proteins enriched on construction **2** (i.e. antiparallel topology with two lateral loops). KEGG analysis of Common group proteins (**1a** and **2**) and proteins enriched on construction **2** showed similar significant enriched pathways compared to those covered by total constrained G4 interacting proteins (see Figure 2A).

**Supplementary Figure S5.**
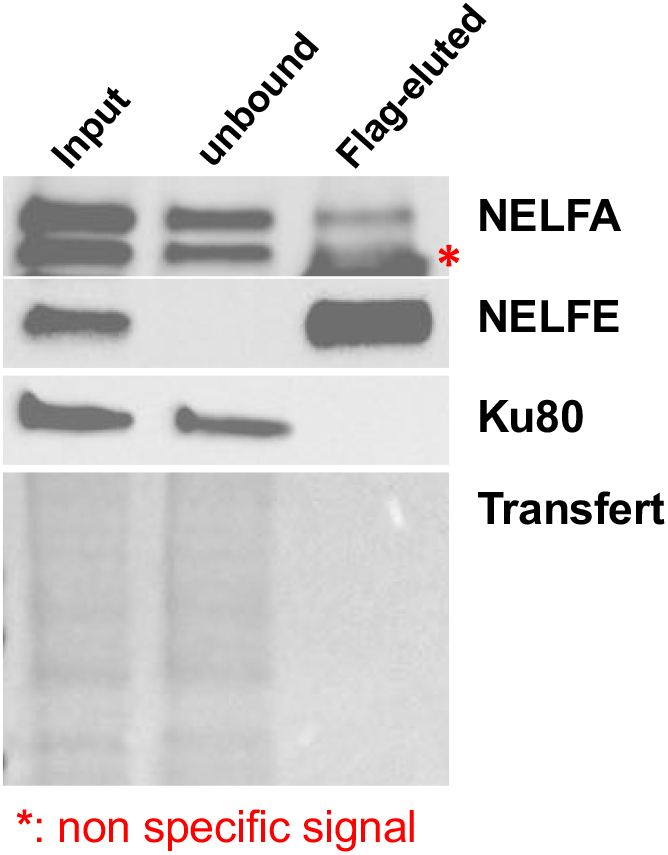
Western-blot detection of NELFE and NELFA proteins immunoprecipitated from a HeLa cell line overexpressing an ectopic Flag-tagged form of the NELF-E protein (Flag-eluted material).

